# Design and analysis of CRISPR-based underdominance toxin-antidote gene drives

**DOI:** 10.1101/861435

**Authors:** Jackson Champer, Samuel E. Champer, Isabel Kim, Andrew G. Clark, Philipp W. Messer

**Affiliations:** Department of Computational Biology, Cornell University, Ithaca, NY 14853; Department of Molecular Biology and Genetics, Cornell University, Ithaca, NY 14853

## Abstract

CRISPR gene drive systems offer a mechanism for transmitting a desirable transgene throughout a population for purposes ranging from vector-borne disease control to invasive species suppression. In this simulation study, we assess the performance of several CRISPR-based underdominance gene drive constructs employing toxin-antidote principles. These drives disrupt the wild-type version of an essential gene using a CRISPR nuclease (the toxin) while simultaneously carrying a recoded version of the gene (the antidote). Drives of this nature allow for releases that could be potentially confined to a desired geographic location. This is because such drives have a nonzero invasion threshold frequency, referring to the critical frequency required for the drive to spread through the population. We model drives which target essential genes that are either haplosufficient or haplolethal, using nuclease promoters with expression restricted to the germline, promoters that additionally result in cleavage activity in the early embryo from maternal deposition, and promoters that have ubiquitous somatic expression. We also study several possible drive architectures, considering both “same-site” and “distant-site” systems, as well as several reciprocally targeting drives. Together, these drive variants provide a wide range of invasion threshold frequencies and options for both population modification and suppression. Our results suggest that CRISPR toxin-antidote underdominance drive systems could allow for the design of highly flexible and potentially confinable gene drive strategies.

## INTRODUCTION

It is possible to engineer alleles that can spread at a higher rate than expected under Mendelian inheritance by biasing their own rate of transmission. These are known as “gene drives”^1–7^. While there are many examples for naturally occurring alleles with super-Mendelian transmission, purpose-engineered gene drive systems are gaining increasing interest as methods to spread desirable transgenic packages throughout a species of interest. The potential applications of such drives fall broadly into two categories: those designed for population modification and those designed for population suppression^1–3,5^. A population modification system may, for example, be designed to alter a trait in mosquitoes that results in reduced transmission of certain diseases. Another example of a modification system would be one designed to help rescue an endangered species from a disease that is spreading too rapidly for natural evolutionary processes to counter^1–3^. Suppression gene drives, on the other hand, could be intended to result in the eradication of an invasive species, an agricultural pest, or a disease vector^1–3,5^.

CRISPR homing drives have recently been the subject of intensive research efforts. These drives utilize CRISPR-Cas9 to cleave the wild-type allele in heterozygotes at a guide RNA (gRNA) target site. The severed DNA end is then repaired by homology-directed repair during which the cell uses the drive-carrying chromosome as a template, resulting in a cell homozygous for the drive allele. This process is intended to occur in germline cells, so that drive carriers will pass the drive on to their offspring at an increased rate. Homing drives have already been developed in a variety of species^8–12^. While resistance alleles have proven a substantial obstacle for these drives, two studies have recently demonstrated a successful modification drive in flies^13^ and a successful suppression drive in mosquitoes^14^.

One problematic feature of homing-type systems is that they are so-called “global” drives. Unless they have a very high fitness cost, the introduction of even a small number of drive-carrying individuals will likely result in the drive spreading throughout an entire population^15^. For target species such as mosquitoes that are capable of long-distance migration by piggybacking on human-based modes of travel as well other means such as high-altitude air currents, this implies that a single release of a homing drive could potentially result in the drive spreading throughout the entire range of the species. In some scenarios, this may be desirable, such as when the drive in question is engineered for disease prevention and when there exists a high degree of social approval. However, this perhaps limits this strategy to a few especially harmful species, such as disease-carrying ticks and mosquitoes. For many other potential applications, confinement to a target area would be highly desirable.

Several designs have already been proposed for gene drives that could potentially be confined. These are generally based on the concept of an invasion threshold, referring to drives that only tend to spread when introduced above a certain threshold frequency, while being removed from the population when introduced below this frequency. Examples of such drives include the *Medea* toxin-antidote (TA) system^16^, variations thereof^17^, *Wolbachia* TA elements^18^, reciprocal chromosomal translocations^19^, and a single locus underdominance TA system called RPM-Drive^20,21^. These^22–34^ and other^35–41^ TA designs have been modeled computationally. Yet both demonstrated and proposed systems have proven difficult to successfully engineer in species of interest due to the need for highly specific targets, promoters and RNAi elements, methods that tend to introduce high fitness costs, and other factors. All proposed drives with invasion thresholds thus far have focused on population modification strategies, while there has not yet been a proposed design for a threshold-dependent suppression drive.

CRISPR toxin-antidote gene drives have recently been devised as an alternative class of drive systems that are substantially less vulnerable to resistance than CRISPR homing drives and often promise easier construction as compared to RNAi-based systems^42–44^. Several of the suggested designs also have invasion thresholds, meaning that they are unlikely to spread through a population unless the drive allele is present above a critical frequency. In a CRISPR-based TA system the “toxin” is a Cas9 element with gRNAs programmed to cut an essential gene on the wild-type chromosome, where cleavage-repair will typically result in a disrupted version of the target gene. The “antidote” element is a functioning copy of the target gene that is located within the drive allele and is recoded to no longer match the drive’s gRNAs so that it is not subject to disruption by the drive. Thus, individuals who inherit only a toxin-disrupted allele suffer from a toxic effect, while individuals who only inherit the drive, or who inherit both a disrupted allele as well as the antidote, do not experience the deleterious toxic effect. By this mechanism, the relative frequency of the drive should increase over time as wild type-alleles are removed from the population.

Two such CRISPR-based TA systems, termed TARE (Toxin-Antidote Recessive Embryo drive) and ClvR (Cleave and Rescue), have already been demonstrated in *Drosophila* – both were able to rapidly spread through cage populations^43,44^. In a recent study, we have modelled several additional designs by varying the nature of the target gene and the expression profile of the drive promoter, showing that these designs can in principle be used for both population modification and suppression strategies^42^. However, most of these designs were so-called “regional” drives, which have a nonzero introduction threshold only when the drive allele carries fitness costs in addition to those imposed by the drive mechanism itself (*i.e.*, costs associated with the disruption of wild-type alleles). While it seems a reasonable assumption that any gene-drive system should have at least some such additional fitness cost (due to expression of the large Cas9 protein, for example), the thresholds of these systems may still be too low for applications where more stringent confinement is desired.

In this study, we propose and model several new designs for CRISPR-based TA systems that can be configured to function as “local” drives by employing underdominance principles. Such local drives are characterized by nonzero invasion threshold frequencies even without additional fitness costs, thereby offering a higher degree of confinement than regional drives. We will present a selection of several such systems, focusing on variants with particularly unique or interesting properties and which seem plausible to construct with current technology. These drives feature a wide span of invasion thresholds and include both population modification and suppression drives.

## METHODS

### Population model

Our simulation model considers a single panmictic population of sexually reproducing diploids with non-overlapping generations. Each individual is specified by its genotype at the drive locus (or drive loci for strategies involving more than one locus) and any additional potential drive target loci, if different from the drive loci.

The fitness of an individual is influenced by its genotype, as defined for the specific drive strategy. Note that there are two different levels at which selection operates in our model. The first level constitutes the drive mechanism itself (based on the disruption of wild-type alleles). This may result in some individuals being sterile or some offspring being nonviable. The second level constitutes any additional fitness costs directly imposed by drive alleles (for example due to expression of the Cas9 protein, the presence of certain other drive elements, or a potential payload). When we refer to the *fitness* (*ω*) of an individual, we always refer to this second level. This fitness then affects mating success in males and fecundity in females, according to the rules described below.

In any given generation, the state of the population is defined by the numbers of male and female adults of each genotype. To determine the state of the population in the next generation, each female first selects a random candidate among all males in the population. The candidate is then accepted for mating at a rate equal to his fitness value (e.g. a male with fitness 0.5 would be selected half the time). If the candidate is rejected, the female chooses another random candidate; if the female rejects ten candidates, she does not reproduce. After a mate has been selected, we set the fecundity of the female (i.e. her expected number of offspring) to twice her fitness value, multiplied by a density dependent scaling factor *σ* = *β* / ((*β* − 1) * *N* / *K* + 1). Here, *β* is the maximum low-density growth rate, *N* is the current population size, and *K* is the carrying capacity of the population. As default values for our model, we used *β* = 10 and *K* = 100,000. A *β* value of 10 means that the population is able to experience a 10-fold growth rate per generation at very low density. The actual number of offspring for the female is then determined by a draw from a binomial distribution with 50 trials and *p* = *ω* * *σ* / 25, yielding a maximum of 50 offspring and an average of 2 offspring for a female with fitness *ω* = 1.0 in a population at carrying capacity (before accounting for potential sterility or nonviability of offspring due to the drive mechanism). These parameters are expected to result in logistic growth dynamics where the population is pushed towards carrying capacity when disturbed, except in the presence of a sufficiently robust suppression drive system.

Each offspring generated this way is assigned a random sex, and its genotype is determined by randomly selecting one allele from each parent, with the possibility that wild-type alleles may be converted into disrupted alleles by drive activity. Each of the parents’ drives disrupts its target site at a rate corresponding to the germline cut rate of the promoter. Next, additional disruption can occur in the embryo, where each of the mother’s drives disrupts the wild-type allele at a rate equal to the embryo cut rate of the promoter. Offspring with nonviable genotypes are then removed.

We generally assume in our model that the additional fitness costs carried by a drive are multiplicative: if drive homozygotes have a fitness *ω*, drive heterozygotes will have fitness 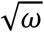. For purposes of easily comparing the drives, in drive systems that consist of multiple drive alleles at different loci, only alleles at one drive locus are allowed to carry additional fitness costs (in suppression drives, if one type of drive allele disrupts a fertility gene with its presence, this is the allele which has the fitness cost). Unless otherwise specified, idealized drives (line graphs) are assumed to carry no fitness cost, and nonidealized drives (heatmaps) are assumed to have *ω* = 0.95.

At the outset of the simulations, a percentage of individuals in the simulation are set as carriers of the selected gene drive system (for modification drives, individuals are set to be homozygous; for suppression drives, individuals are set to be heterozygous). The simulation is then run for 100 generations, with several metrics of drive performance tracked. All models were implemented in the SLiM simulation framework^45^.

### Data output

At each generational step in our simulations, we record the frequency of the drive allele(s) in the population, the percentage of individuals that are drive carriers, the population size, and the genetic load of the drive on the population. The genetic load that the drive imposes on the population in a given generation is defined as 1 − *N*_*act*_/*N*_*exp*_, where *N*_*act*_ is the actual number of individuals observed in the next generation and *N*_*exp*_ is the number expected if all individuals were wild type (according to our logistic growth model). By taking the average of the genetic load over several generations after the drive has reached fixation or equilibrium, we can assess the reproductive burden that a drive imposes on a population.

We used a numerical approach to estimate the required introduction threshold of a drive. In particular, we considered a given introduction frequency to be above the threshold if the drive increased in frequency during any ten consecutive generations within the fifty generations following the introduction of the drive. The final thresholds assigned to the drives are the lowest introduction rates where, in half or more simulations, this condition was satisfied.

### Data generation and software

Simulations were run on the computing cluster of the Department of Computational Biology at Cornell University. All simulations were run using SLiM version 3.3. Data processing, analysis, and figure preparation were performed in Python 3.7.4. The SLiM program and parameter files allowing the reader to reproduce all simulations presented here are available on GitHub (https://github.com/MesserLab/TA-Underdominance-Drives).

## RESULTS

### General overview of underdominance TA systems

All CRISPR-based TA systems work via the same fundamental strategy: the drive disrupts an essential gene while also providing a “rescue” element for the gene. This results in removal of wild-type alleles, allowing the drive to increase in relative frequency over time. However, a multitude of possible configurations for such drives are conceivable by varying the type of target gene, the expression profile of the endonuclease, the genomic location of the drive, etc. Several of these have been modeled previously^42–44^, but so far, all of these studies have focused on systems with low invasion threshold frequencies. Here, we focus on CRISPR TA systems that achieve higher thresholds by use of underdominance.

The underdominance principle is that drive/wild-type heterozygotes are less successful than either drive or wild-type homozygotes. This generally results in a nonzero invasion threshold frequency for the drive even when it does not carry any additional fitness costs (note that whenever we refer to the fitness costs of a drive, we mean additional costs that do not result from the drive mechanism itself, i.e. the disruption of wild-type alleles). The invasion threshold represents the critical frequency above which drive-carrying individuals (homozygotes for modification drives and heterozygotes for suppression drives) need to be introduced to the population so that the drive is expected to increase in frequency (either to fixation or some equilibrium frequency). When introduced below this frequency, the drive is expected to be lost from the population. Such drives with nonzero invasion thresholds even in the absence of fitness costs are often termed “local” drives, though their exact degree of localization depends on the threshold and numerous other ecological factors. This is in contrast to previously considered “regional” CRISPR TA systems that do not employ underdominance mechanisms and generally have a nonzero invasion threshold frequency only if the drive has additional fitness costs^42–44^, or “global” TA systems that have a zero invasion threshold even under a wide range of fitness costs^42^. In a scenario where two demes are linked by migration, and assuming panmixia within each deme, local and (to lesser extent) regional drives can be successfully confined to one of the demes if the migration rate between them is below a critical threshold^22,23,29,32,41,46^. This migration threshold, of course, depends on the invasion threshold frequency but also accounts for the fact that even a low rate of migration could eventually lead to the drive exceeding its invasion threshold in the other deme when drive alleles can accumulate over time^22,23,29,32,41,46^.

In this study, we propose several novel configurations for underdominance CRISPR TA drives, including strategies for both population modification and suppression. We first discuss the general components of these drives, focusing specifically on the nuclease promoter, the type of target gene and rescue element, and the different architectures in which these elements can be arranged. This is followed by a consideration of specific drive systems and computational analysis of their expected performance through simulations in a simple panmictic population model (see Methods). While the precise dynamics of these drives in a real-world application would undoubtedly depend on various aspects of the ecology of the target species, our goal here is to introduce the general principles and performance characteristics of these drives as a necessary starting point for future experimental work and more detailed modeling studies.

### Components of TA systems

#### Nuclease promoter

The choice of promoter regulates the expression of the Cas9 (or another nuclease) and thus determines the rate and timing at which the drive disrupts wild-type target genes. The choice of promotor can thereby affect whether cutting occurs primarily in the germline or also during early embryo development due to maternally deposited Cas9. An idealized (in line graphs) germline promoter (G promoter) would result in 100% cutting activity in the germline and no activity in the early embryo from maternally deposited Cas9, while an idealized germline and embryo promoter (GE promoter) would have 100% cutting activity at both stages. For nonidealized forms (in the heatmaps), we assume in our model that germline cut rate is 99% in both cases, as seems to be the case with most experimentally tested drive promoters^47,48^, while embryo activity is 5% for a germline-restricted promoter^14,48^ and 95% for a promoter that also has high activity in the early embryo^49,50^.

A promoter with somatic expression^47,49^ (GES promoter) additionally induces Cas9 expression in non-germline cells, leading to the formation of disrupted wild-type target alleles if a drive allele is present. We model such promoters as having activity in the germline and embryo as above, but individuals with drive alleles and wild-type target sites are then considered to also have disrupted target sites when determining viability and fitness in our simulation model. This represents a GES promoter that induces substantial somatic Cas9 expression.

#### Target gene and rescue element

The target of a TA system can be either dominant or recessive. In TA systems with dominant targets (e.g. TA**D**E, Toxin-Antidote Dominant Embryo drive), an individual is nonviable if it has less than two functioning copies of the gene, unless drive rescue provides the equivalent thereto. If the drive target is recessive (e.g. TA**R**E, Toxin-Antidote Recessive Embryo drive), the individual is fully viable if it has at least one functioning copy of the gene, or the equivalent provided by drive rescue. “Regional” TARE and TADE drives (including a suppression form of the latter) have been modeled previously with both G and GE promoters^42–44^, but in this study, we present additional unique configurations of drives with TARE and TADE elements.

The rescue provided by the drive could be equivalent to the functionality of the wild-type allele, or, by using different regulatory sequences or otherwise modifying the rescue portion of the drive, could be configured to provide half the functionality of the wild-type allele (e.g. TA**H**RE, Toxin-Antidote Half-rescue Recessive Embryo drive – two copies of the drive are equivalent to a single wild-type copy of the target gene). Similarly, by using two rescue gene copies in a single drive allele, the drive could provide double functionality (e.g. TA**D**DE, Toxin-Antidote Double-rescue Dominant Embryo drive – one drive allele is equivalent to two copies of the wild-type target allele). TADDE drive has been modeled previously in a simple configuration that resulted in a “regional” drive^29^.

Four meaningful permutations of these systems can be envisioned: double rescue with a haplolethal target (TADDE), single rescue with a haplolethal target (TADE), single rescue with a haplosufficient target (TARE), and half rescue with a haplosufficient target (TAHRE). Generally, TARE and TADDE systems are suitable for population modification but lack the ability to impose a substantial genetic load on a population for suppression^42^. TADE^42^ and TAHRE systems can optionally introduce strong genetic loads for population suppression (as we will show below), though suppression systems tend to act more slowly than modification systems.

#### Drive architecture

A drive allele can be a “same-site” allele where the drive is co-located with the gene that it provides rescue to, or it can be “distant-site” where the drive provides rescue for a distant gene. Some types of distant-site drives are able to strongly suppress a population if the drive is located in an essential but haplosufficient female fertility gene (or male-specific, and/or affecting viability instead of fertility) and disrupts the gene with its presence. A female with two copies of the drive is therefore sterile. Both same-site and distant-site alleles can also be used for population suppression if additional gRNAs that target the female fertility gene are included in the drive (without a recoded rescue element), even if the drive itself is not located in the fertility gene.

Drive arrangements can further be classified into three types: single locus systems with a single drive allele (all previously considered CRISPR TA systems used such a configuration^42–44^), single locus systems with two types of drive alleles, and multi-locus (usually two) systems with a different type of drive allele at each site. In the single-locus system with a single drive allele, the drive targets and provides rescue for the same gene. In the other two systems, the first drive allele targets the gene that the second drive allele provides rescue for, and vice versa.

### Design and analysis of underdominance TA systems

#### 1-Locus 2-Drive TARE

We first propose a system that comprises two TARE drive alleles that occupy the same genetic locus (Figure 1A). A similar RNAi-based system has an invasion threshold frequency of 67%^22,23,25,30,46^. Each of the two drives targets an essential but haplosufficient gene. The first drive targets the gene which the second provides rescue for (via a recoded copy that cannot be targeted by the drive), and vice versa. One of these drive alleles could be in a same-site configuration, or both could be in distant-site configurations. We specifically model the latter system, along with a promoter that yields high Cas9 cutting activity in both the germline and in the early embryo from maternally deposited Cas9, which should be most efficient for standard TARE drives^42^. If the promoter also has somatic activity, then the dynamics of this system should be similar to the 1-locus 2-drive RNAi-based system that has been modeled previously^22,23,25,30,46^ (this system has a slightly higher introduction threshold than the CRISPR 1-locus 2-Drive TARE system).

**Figure 1.**
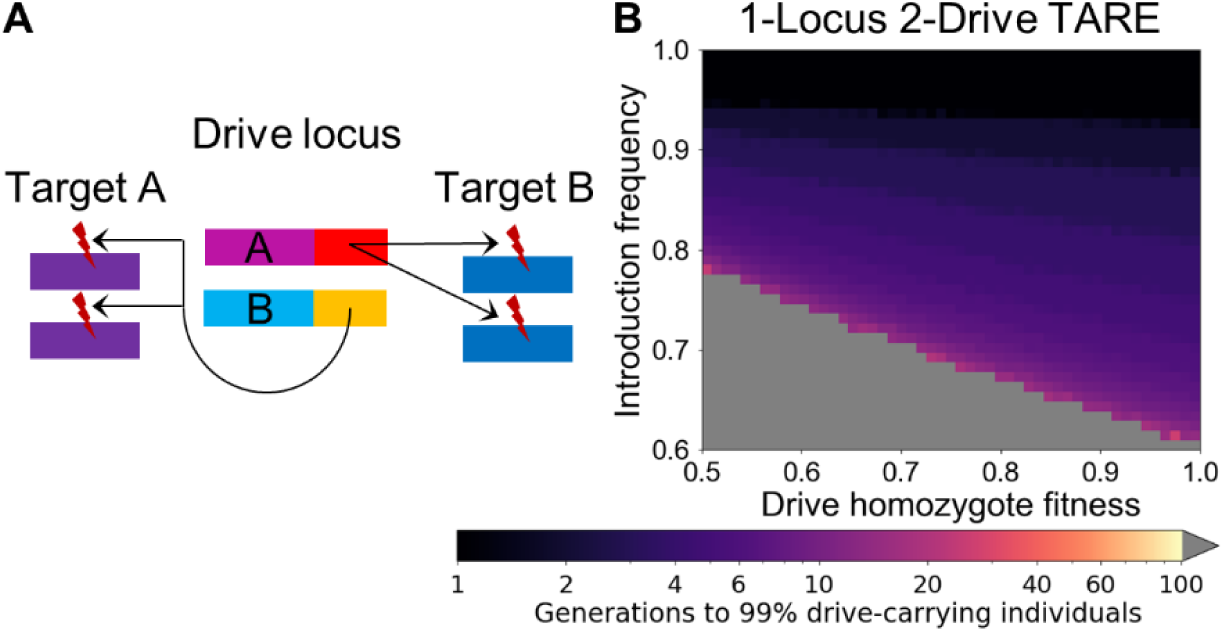
1-Locus 2-Drive TARE. (**A**) In the 1-Locus 2-Drive system, two TARE drive alleles (A and B) are situated at the same locus. Each targets a different essential but haplosufficient gene while providing rescue for the other drive allele’s target. (**B**) The time at which a TARE-based drive with a GE promoter is expected to reach 99% of individuals in the population in our simulation model with varying drive-carrying individual introduction frequency and drive fitness. Released individuals have one copy of each drive allele at the drive site. Grey indicates that the drive was eliminated within 100 generations.

The reciprocal targeting scheme of the drives, along with the fact that both drive alleles are situated at the same genomic locus, means that after all wild-type alleles in a population have been disrupted, the only viable genotype is one in which both drive alleles are present. Furthermore, all crosses involving females with a copy of each drive will result in some offspring not surviving (half in crosses with males with one copy of each drive, and all in crosses with wild-type males). Thus, after fixation, this drive imposes a genetic load of 0.5 on the population even if the drive carries no additional fitness cost (see Methods). This will likely result in a modest suppressive effect if the drive alleles fixate, depending on the ecological characteristics of the organism in question. Because of these dynamics, this drive has the highest threshold of all the modification drives we analyze in this study. With a GE promoter, the drive requires an introduction above 61% to spread in the absence of additional fitness cost (63% for a G promoter). However, the drive can remove wild-type alleles quickly when present at such high frequencies, resulting in the combined drive allele frequency rapidly reaching 100% if released above its threshold (Figure 1B).

#### 2-locus drives

We next propose a set of drives that comprise TARE, TADE, and TADDE alleles at two genetic loci, with each locus containing a different drive allele (thus allowing any or all the drives to be same-site). Such a system based on RNAi elements has already been well-studied, with an invasion threshold of 27% in the absence of fitness costs^22,23,25,27,28,30,33,46^. As in the 1-locus 2-drive system, each drive allele targets the gene for which the other provides rescue (Figure 2A). However, because each drive allele has its own locus, drive alleles are removed at a lower rate than wild-type alleles in more crosses, substantially decreasing invasion threshold frequencies compared to the 1-locus drive arrangements. For example, if both drives are TARE type with GE promoters, crosses between wild-type males with heterozygous (at both loci) females result in 3/4 of offspring being non-viable, removing wild-type and drive alleles at a 5:1 ratio. Crosses between two heterozygotes result in 7/16 of offspring being nonviable, removing wild-type and drive alleles at a 5:2 ratio. Because of the relatively high rate of wild-type allele removal, this drive has only a modest invasion threshold of 18% in the absence of fitness costs. The drive rapidly reaches all individuals (Figure 2B), but as in a single-locus TARE drive, it takes longer for drive alleles to fixate, and the drive frequency will instead reach an equilibrium if drive alleles have a fitness cost^42^. The drive has similar performance with a somatic GES promoter but is somewhat slowed if a germline-only promoter is used (though in the latter case, the invasion threshold is also slightly reduced to 17%).

**Figure 2.**
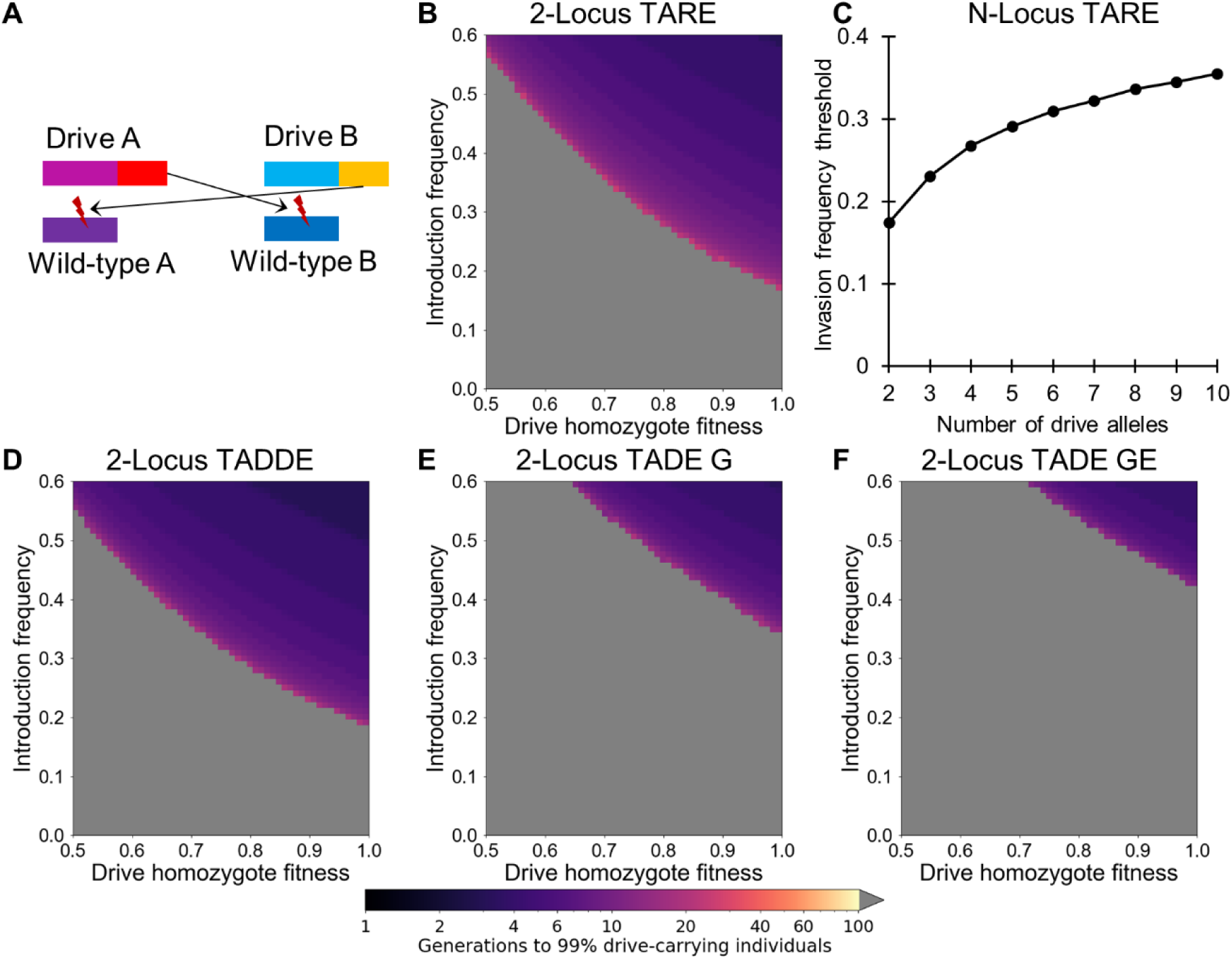
2-Locus drives. (**A**) In the 2-Locus drive systems, two drive alleles (both providing “same-site” rescue in this example) each target an essential but haplosufficient gene while providing rescue for the other drive allele’s target. (**B**) The time at which a 2-locus drive with unlinked, same-site TARE alleles with a GE promoter is expected to reach 99% of individuals in the population with varying introduction frequency and drive fitness. (**C**) The introduction frequency thresholds for TARE drives with additional loci, where each drive cyclically provides rescue for the target of the previous drive. (**D**) As in (B), but for TADDE alleles. (**E**) As in (B), but for TADE alleles with a germline (G) promoter. (**F**) As in (B), but for TADE alleles with a promoter causing cutting activity in the germline of both sexes and in embryos of drive-carrying females (GE). Released individuals are homozygous for all drive alleles. Grey indicates that the drive was eliminated within 100 generations.

A drive system can also be implemented with more than just two loci and types of drive alleles. In this case, each drive targets the gene that the next drive provides rescue for, with the last drive targeting the first. Figure 2C shows the invasion threshold frequencies of such systems based on TARE alleles. As the number of drives increases, the number of drive alleles removed by mutual disruption increases, thus increasing the threshold frequency.

In a 2-locus TADDE drive, the doubled rescue element results in unchanged performance when using a promoter with or without embryo activity, or even a promoter with somatic expression in the offspring. This drive spreads slightly faster than the 2-locus TARE drive since wild-type alleles are removed more quickly (Figure 2D). Crosses between a drive/wild type heterozygote at both loci with a wild-type individual result in 3/4 of offspring being nonviable, representing a removal of wild-type and drive alleles at a ratio of 5:1. Crosses between two heterozygotes at both loci results in 7/16 of offspring being nonviable and represents a removal of wild-type and drive alleles at a 5:2 ratio. These dynamics result in an invasion threshold of 19% in the absence of drive fitness cost, slightly higher than the TARE version.

A 2-locus TADE system has substantially different performance with a G (Figure 2E) or GE (Figure 2F) promoter. With a G promoter, all crosses involving homozygotes have a regular number of offspring. However, with a GE promoter, crosses between any drive-carrying female and a wild-type male yield no offspring. A cross between a heterozygote female and a wild-type male, when the drive utilizes a G promoter, results in 3/4 of offspring being non-viable, representing a removal of wild-type and drive alleles at a 5:1 ratio. A cross of heterozygotes at both loci results in only 1/16 of offspring being viable, representing a removal of wild-type and drive alleles from the population at an 8:7 ratio. These drive dynamics result in an invasion threshold of 33% when using a G promoter and 43% when using a GE promoter.

#### 2-locus TADE suppression

We next propose a 2-locus TADE system wherein one of the drive alleles is placed in a female fertility gene (Figure 3A). This drive should be able to induce a high genetic load on a population, resulting in effective population suppression (Figure S1). In an alternative construction, a drive allele could instead target a female fertility gene with additional gRNAs (without providing rescue) and have similar suppressive performance. For suppression drives, we model heterozygote releases, since heterozygotes may be easier to generate and are fully fertile (the system technically has better performance if released males are homozygous and released females are heterozygous for the suppression element and homozygous for the second). As with the modification system, the drive system works best with a G promoter (Figure 3B) and has a higher invasion threshold and slower spread with a GE promoter (Figure 3C). In both cases, the invasion thresholds for the suppression variant are substantially higher than for the modification variant. Due to the many drive alleles that are in nonviable or sterile individuals, the drive has a relatively high invasion threshold of 83% with a G promoter and 88% with a GE promoter. The application of these systems would therefore require an isolated area and very large release sizes.

**Figure 3.**
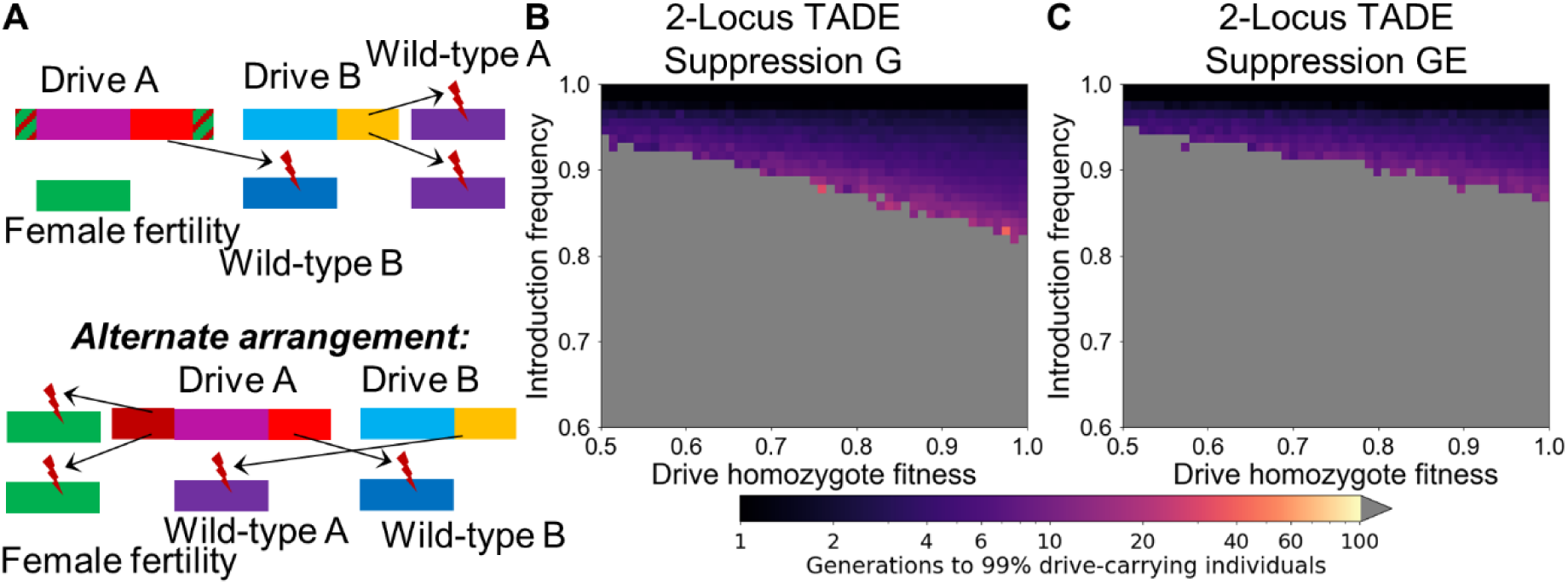
2-Locus TADE suppression drives. (**A**) A 2-locus TADE drive will function as a suppression drive if one of its drive alleles is “distant-site” and located in an essential but haplosufficient female fertility gene (or any single-sex fertility or viability gene), disrupting the gene with its presence. Alternatively, the drive allele can simply target the fertility gene with additional gRNAs, allowing a “same-site” arrangement. (**B**) The time at which a 2-locus TADE suppression drive (with one allele placed in a female fertility gene and the other allele in a same-site configuration, with all components genetically unlinked) with a germline promoter (G) is expected to reach 99% of individuals in the population with varying introduction frequency and drive fitness. (**C**) As in (B), but for TADE alleles with a promoter leading to cutting activity in the germline of both sexes and in embryos of drive-carrying females (GE). Released individuals are heterozygous for all drive alleles. Grey indicates that the drive was eliminated within 100 generations.

#### TADE underdominance

We next propose a TADE underdominance system. This drive is similar to the previously considered TADE drive^42^, but uses a GE promoter (Figure 4A). Such a drive is expected to exhibit underdominance characteristics as well as a threshold. Here, we consider a same-site system, but a similar system based on a distant-site location with a GE promoter has been modeled previously and has highly similar characteristics^44^. We also consider a variant of the drive with a promoter that exhibits ubiquitous Cas9 somatic activity.

**Figure 4.**
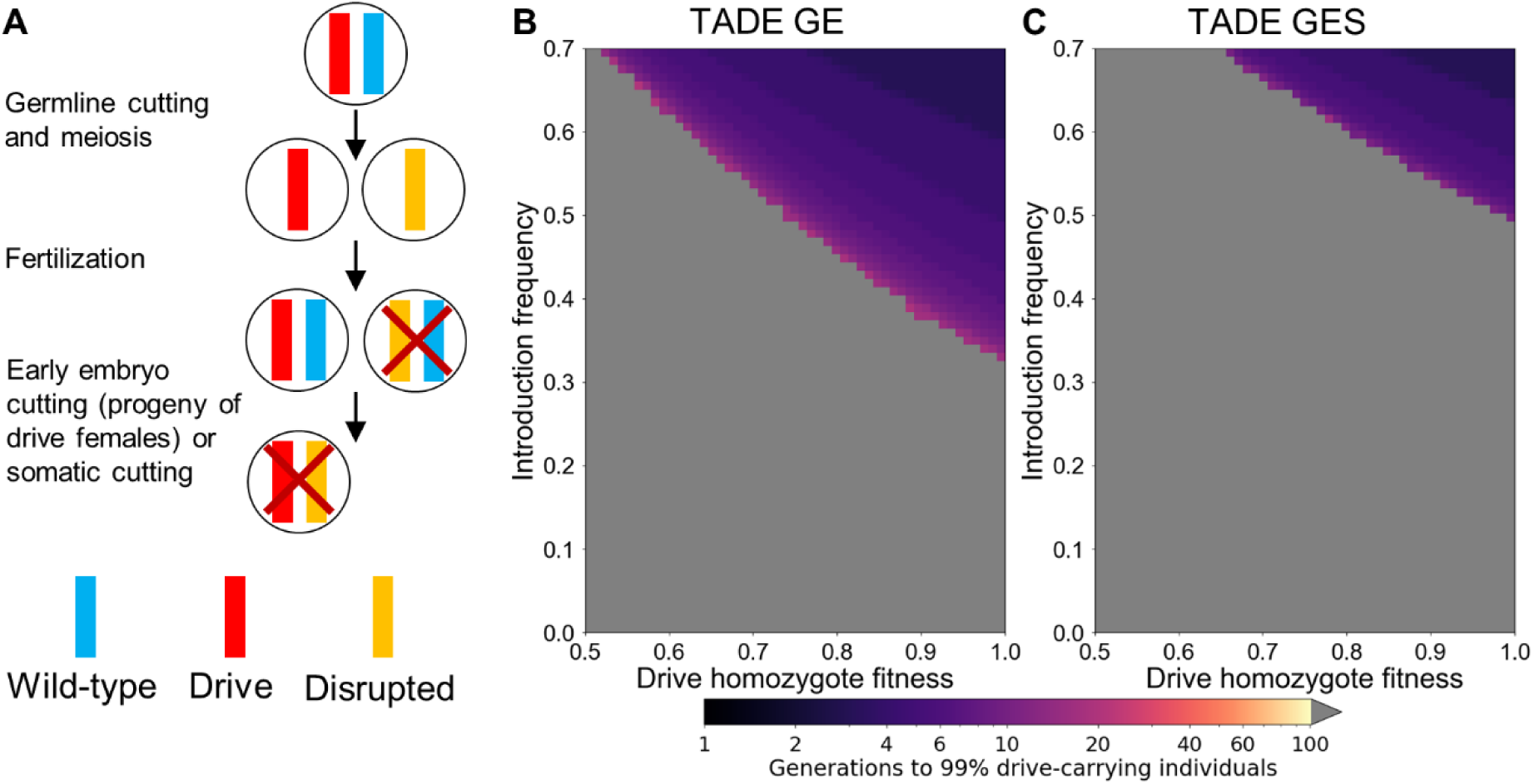
TADE underdominance drives. (**A**) A TADE drive acts as an underdominance drive if it utilizes a promoter that induces cutting activity in the germline and early embryo (GE) due to maternally deposited Cas9 and gRNA. The promoter can also have leaky somatic expression (GES), resulting in cutting of wild-type alleles in somatic tissues by the drive. (**B**) The time at which a same-site TADE drive with a GE promoter is expected to reach 99% of individuals in the population with varying introduction frequency and drive fitness. (**C**) As in (B), but for TADE alleles with a promoter that also drives somatic cutting (when this takes place in our model, such individuals are all nonviable). Released individuals are homozygous for the drive allele. Grey indicates that the drive was eliminated within 100 generations.

As implemented with a GE promoter, offspring from crosses between a drive homozygous male and wild type female are all viable, while crosses between a drive-carrying female and a wild type male produce no offspring. A male heterozygote crossed with a wild type female results in half the offspring being viable, removing only wild type alleles from the population. A cross between heterozygotes results in 3/4 of potential offspring being nonviable, producing only homozygotes and representing a removal of wild-type and drive alleles at a 2:1 ratio. This drive has a moderate introduction threshold of 1/3 in the absence of allelic fitness costs, and above this threshold it spreads rapidly to fixation (Figure 4B).

When a TADE underdominance drive uses a GES promoter that also exhibits somatic activity, heterozygotes are nonviable, and crosses between drive and wild-type individuals are assumed to never produce viable offspring. Therefore, in the absence of fitness costs, the drive has a 50% invasion threshold, but it still spreads rapidly when released above its threshold (Figure 4C). Such a system would have highly similar dynamics to single-locus single-allele RNAi based systems^26,29,32^.

#### TADE underdominance suppression

We next propose a TADE underdominance suppression drive. This drive can take the form of a distant-site TADE underdominance drive with a GE promoter located inside a haplosufficient but essential female fertility gene (or other suitable location), disrupting the gene with its presence (Figure 5A). Alternatively, the drive can be same-site with the viability target and have additional gRNAs targeting the female fertility gene without rescue (Figure 5A). These two architectures were found to have similar performance. Because of the relatively many individuals that are nonviable or sterile from the effects of the drive, TADE underdominance suppression has a high invasion threshold frequency of 61% in the absence of drive fitness cost (Figure 5B).

**Figure 5.**
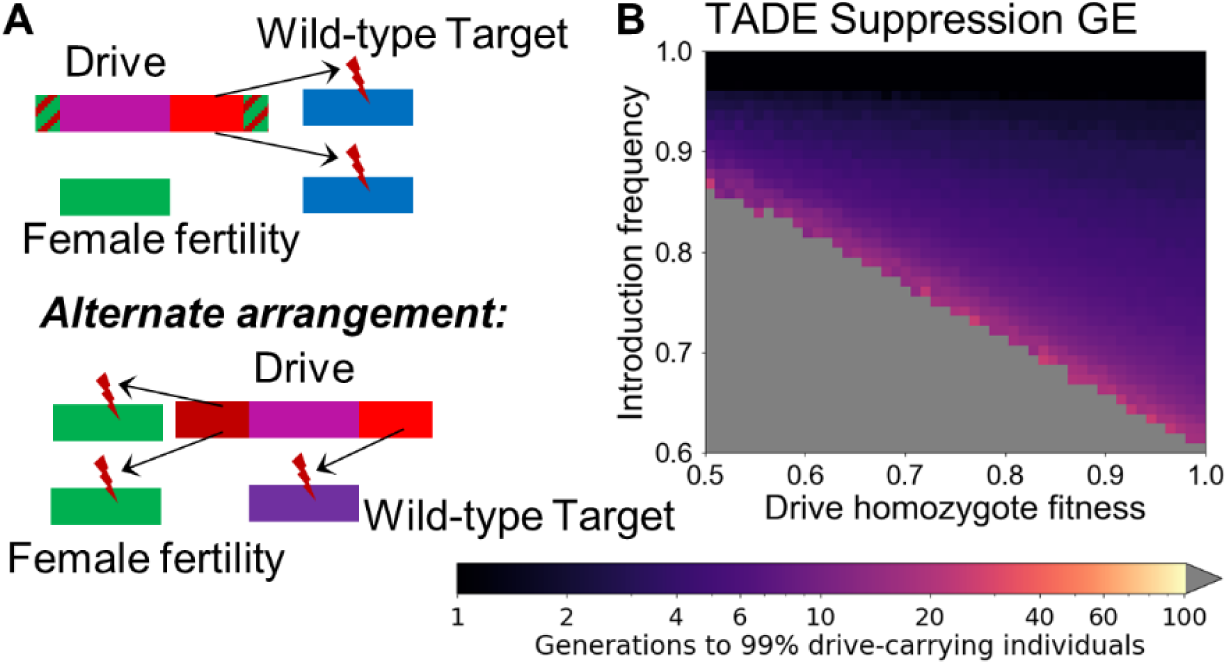
TADE underdominance suppression drive. (**A**) A TADE underdominance drive (with a GE promoter for activity in the germline and early embryo if the mother had a drive) will function as a suppression drive if one of its drive alleles is “distant-site” and located in an essential but haplosufficient female fertility (or any single-sex fertility or viability gene), disrupting the gene with its presence. Alternatively, the drive allele can simply target the fertility gene with additional gRNAs, allowing a “same-site” arrangement. (**B**) The time at which a TADE underdominance suppression drive (placed in a female fertility gene) is expected to reach 99% of individuals in the population with varying introduction frequency and drive fitness. Released individuals are heterozygous for the drive allele. Grey indicates that the drive was eliminated within 100 generations.

#### Variable embryo cut rate and target haploinsufficiency

It is possible that a particular promoter may have an intermediate level of cutting activity in the early embryo from maternally deposited Cas9^10,14,47,48,51^. A TADE modification or suppression drive configured with such an intermediate promoter has a lower threshold than one with a GE promoter, with the exact level being dependent on rate of embryo cutting (Figure S2). Drives with these promoters spread rapidly, though less when the introduction frequency is close to their invasion threshold (Figure S3). While this potentially increases the range of invasion thresholds and confinement for a drive if the embryo cut rate can be adjusted, such an approach should be treated with caution. This is because for intermediate levels of embryo cutting, genetic background can significantly impact the cutting rate on an individual-to-individual basis^10,51^, and this could result in a change of the average rate over time as individuals with higher rates tend to be rendered nonviable more often.

Another variant is for the target gene to have an intermediate level of haploinsufficiency, thus placing it between the haplosufficient target of a TARE drive and the haplolethal target of a TADE drive. For a GE promoter, an increase in the degree of haploinsufficiency will steadily increase the invasion threshold from zero to the level of our TADE underdominance drives (Figure S4). We also considered both modification and suppression drives in which both the target haploinsufficiency and the cut rate in the early embryo were allowed to vary (Figure S5). When haploinsufficiency is low, a GE promoter results in faster drive spread, and when haploinsufficiency is high, a G promoter is optimal for maximizing the rate of drive spread. The optimal level of embryo cutting transitions quickly from 100% to 0% at an intermediate level of haploinsufficiency (Figure S5). This is usually around 0.2-0.5, with the exact level of the transition varying based on the specific type of drive (modification vs. suppression) and the initial release frequency. However, the rate of embryo cutting has little effect on the rate of spread of the drive in this intermediate region, as long as the initial release level is above the invasion threshold, which is dependent on both the embryo cut rate and the degree of target haploinsufficiency. Note that as the degree of haploinsufficiency decreases, a suppression drive would induce a lower genetic load on a population^29^, independent of the embryo cut rate.

#### TAHRE modification and suppression drive

We finally propose a TAHRE drive system. This drive targets an essential but haplosufficient gene and provides only half rescue (if no wild-type copies of the target gene are present, two copies of the drive are required for viability). Thus, in contrast to a TARE drive, drive/disrupted target heterozygotes are nonviable (Figure 6A). This system is similar in concept to the previously proposed *Merea* drive based on RNAi^37^. We consider a version of the drive with a GE promoter (Figure 6B).

**Figure 6.**
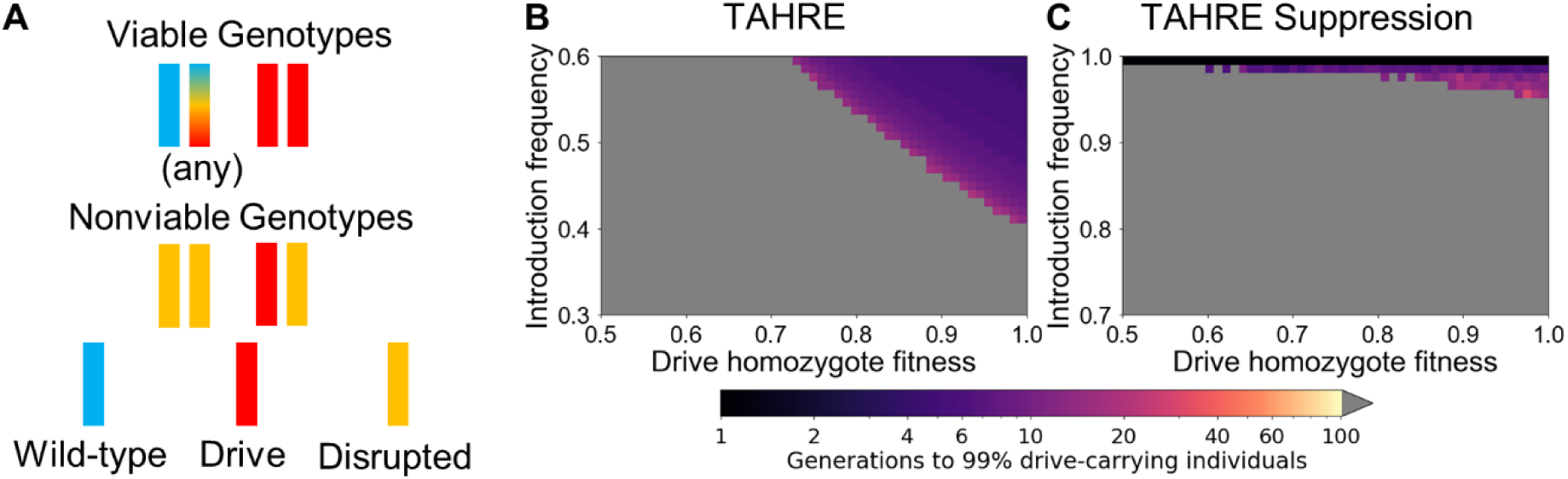
TAHRE drive. (**A**) A TAHRE drive uses a TARE target, but two drive copies are required to provide rescue, thus creating an underdominance system. (**B**) The time at which a same-site TAHRE drive is expected to reach 99% of individuals in the population with varying introduction frequency and drive fitness. Released individuals are homozygous for the drive allele. (**C**) As in (B), but for a TAHRE suppression drive placed in a female fertility gene. Released individuals are heterozygous for the drive allele. Grey indicates that the drive was eliminated within 100 generations.

Because the drive only provides half-rescue, no viable offspring can result from a pairing between a drive-carrying female and a wild-type male. Drive homozygote and heterozygote males produce a regular number of offspring with wild-type females. Crosses between heterozygotes result in 3/4 of offspring being nonviable, representing removal of wild-type and drive alleles at a 2:1 ratio. These characteristics result in the drive requiring a moderate invasion threshold frequency of 41% in the absence of drive fitness cost, which is the same as the *Merea* system^37^ (the invasion threshold is reduced to 35% if a G promoter is used).

When this drive utilizes a promoter that exhibits somatic CRISPR nuclease activity, all drive/wild-type heterozygotes are nonviable, resulting in an invasion threshold frequency of 50% in the absence of drive fitness costs and identical characteristics to a TADE drive with a similar promoter (Figure 4C).

We also propose that a TAHRE drive can be converted to a suppression system in the same manner as described for the TADE drives (Figure 6C). In this case, the drive has a very high invasion threshold frequency of 96% in the absence of fitness costs (the system is nonfunctional with a G or GES promoter). While this threshold is high, such a system is still a gene drive and would theoretically have substantially more suppression power than methods involving sterile insect technique^52^ or releases of *Wolbachia*-carrying males^53^.

#### Target genes with incomplete lethality when disrupted

Thus far, we have considered target genes in which lack of at least one (TARE) or two (TADE) functional wild-type target or drive allele copies must be present to avoid lethality. However, for some possible target genes, instead of a lethal effect, individuals may survive and suffer negative fitness effects. Here, we consider drives with such targets. Instead of nonviability, individuals with insufficient rescue are viable but have their fitness (male mating success and female fecundity, as defined in the methods) multiplied by a fixed value between zero and one. If the drive has a TADE target, the full fitness effect is only suffered by drives lacking any functional wild-type target or drive alleles, and individuals with one such allele have their fitness multiplied by the square root of this fixed value.

In general, drives with incomplete lethality targets have lower invasion thresholds (Figure S6). However, when the fitness of individuals with incomplete rescue is sufficiently high, the construct is no longer able to function as a gene drive (Figure S6). Additionally, higher levels of fitness for individuals with incomplete rescue also slows the spread of the drive (Figure S7) and reduces the potential genetic load that a suppression drive induces in a population when it reaches it final equilibrium frequency (Figure S8).

## DISCUSSION

In this study, we have presented several new CRISPR-based underdominance TA gene drive designs that could allow for the development of localized population modification or suppression drives (Table 1). Such systems can utilize a broad class of target genes and promoters, suggesting that their construction may be feasible in many target organisms. Indeed, two examples of CRISPR-based TA systems have already been demonstrated experimentally^43,44^, and a similar TA underdominance system would likely only require a rearrangement of existing components. Systems using haplolethal targets may be somewhat more difficult to engineer due to the sensitivity of organisms to these gene’s expression levels. They are likely feasible, though, since the targeting of such a gene was recently demonstrated experimentally for a homing drive^13^. On the other hand, it remains unclear how TAHRE rescue elements would be constructed.

**Table 1.** Comparison of CRISPR TA drive types.

| Drive Type | Threshold* | Suppression threshold** | Promoter | Engineering Difficulty |
| --- | --- | --- | --- | --- |
| TARE<sup>42-44</sup> | Low | N/A | Any | Proven<sup>43,44</sup> |
| TADE<sup>42</sup> | Low | Low | G | Medium |
| TADDE<sup>42</sup> | Low | N/A | Any | Medium? |
| TADS<sup>42</sup> | Zero | Zero | Any | High? |
| 1-Locus 2-Drive TARE | High | N/A | Any | Low |
| 2-Locus TARE | Medium | N/A | Any | Low |
| 2-Locus TADE | Medium | Very High | G,GE | Medium |
| 2-Locus TADDE | Medium | N/A | Any | Medium? |
| TADE Underdominance | Medium | High# | GE,GES## | Low? |
| TAHRE | Medium | Very High | G, GE | High? |

\*Thresholds assume a small drive fitness cost and provide an indirect measure for the degree of confinement.
\*\*These are for suppression variants of the drive. N/A indicates that a strong suppression form of the drive is not possible.
#A TADE Underdominance suppression system could have a “medium” threshold if it had intermediate early embryo cutting.

Compared to homing drives, TA underdominance systems allow for local confinement, have lower rates of resistance allele formation by avoiding reliance on the error-prone process of homology-directed repair^42^, and should have a reduced rate of mutational inactivation of potential payload genes, since these are copied only by replication rather than the more error-prone homology-directed repair process. Note, however, that care must still be taken in the construction of such systems to avoid the formation of resistance alleles by undesired homology-directed repair of rescue elements (and not other drive elements). Procedures to mitigate the formation of this type of resistance have already been successfully demonstrated in both same-site^43^ and distant-site^44^ configurations of TA elements. An additional advantage of most TA systems over homing drives is that they often do not require a germline-specific promoter due to their ability to tolerate maternal deposition and subsequent activity of Cas9 in the embryo. Some TA systems can even tolerate ubiquitous Cas9 somatic activity.

The systems presented here are all “local” drives with relatively high invasion threshold frequencies even in the absence of drive fitness costs, spanning a wide range of thresholds (Figure 7). Modeling in continuous space has shown that drives with invasion threshold frequencies at or above 50% in panmictic populations can often fail to persist in well-connected populations unless the drive is released over a wide area, while drives with invasion thresholds below 50% can be invasive in many scenarios^22,34,54^. Thus, the potential for “local” drives with a variety of invasion threshold frequencies (together with previously considered “regional” drives^42–44^) should provide increased flexibility in the development of an appropriate drive for a given application.

**Figure 7.**
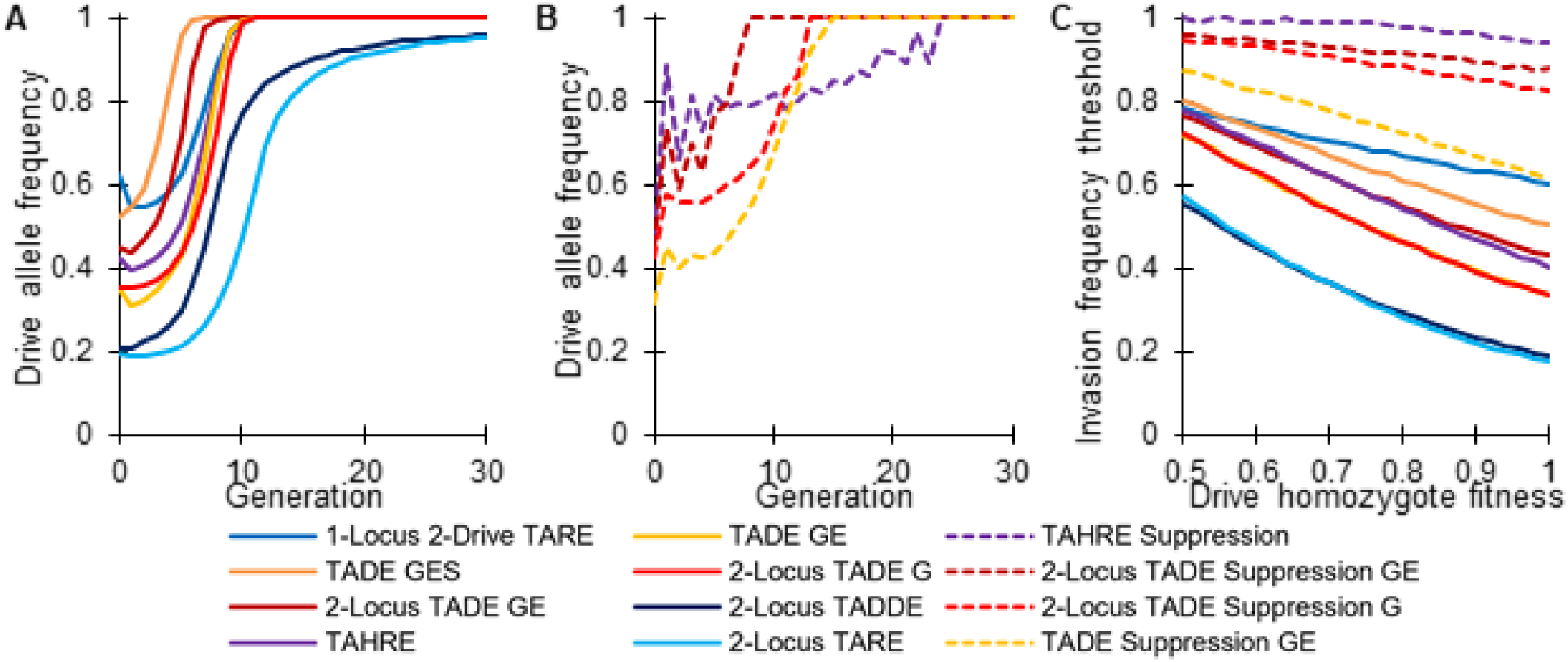
Dynamics of TA underdominance drives. (**A**) Example allele frequency trajectories for modification drives introduced at 2% above their introduction frequency thresholds in our population model. (**B**) Example allele frequency trajectories for suppression drives introduced at 2% above their invasion threshold frequency. (**C**) Invasion threshold frequencies as a function of drive fitness for the different systems. Note that the TADE GE and 2-Locus TADE G drives have the same invasion thresholds. In modification systems, released individuals were homozygous for the drive. In suppression systems, individuals were heterozygous for the drive. G = germline only promoter. GE = promoter with germline and early embryo cutting (in the progeny of drive-carrying females). GES = promoter that induces a high rate of somatic cleavage.

Though we have modelled a wide variety of drives in this study, additional combinations exist that might be suitable in some situations or easier to construct. For example, mutually targeting 2-locus systems can be made up of any combination of individual drive-type elements (TARE, TADE, etc). Suppression drives can utilize additional gRNAs targeting the female fertility gene instead of disrupting the gene by presence of the drive. This could allow a same-site suppression drive to be more easily constructed, with the attendant advantages in rescue element efficiency^43^. Each of these systems could be combined with a tethered homing drive^55^ to provide confinement based on the TA underdominance system, but the power of a homing drive for strong suppression or to spread costly payloads more efficiently.

Overall, our modeling analysis suggests that CRISPR-based underdominance TA systems could be used for both population modification and suppression with a high number of possible variants with different invasion threshold frequencies. Their construction requires elements that have already been demonstrated^43,44^, making the varieties presented here, together with “regional” TA systems^42^, promising candidates for the development of flexible strategies for confined gene drive. Future experimental and computational studies should further characterize underdominance TA drives and assess their implementation in potential target species.

## ACKNOWLEDGEMENTS

This study was supported by New Zealand’s Predator Free 2050 program under award SS/05/01 to PWM, the National Institutes of Health award R01GM127418 to PWM, National Institutes of Health award R21AI130635 to JC, AGC, and PWM, and the National Institutes of Health award F32AI138476 to JC.

## SUPPLEMENTAL INFORMATION

**Figure S1.**
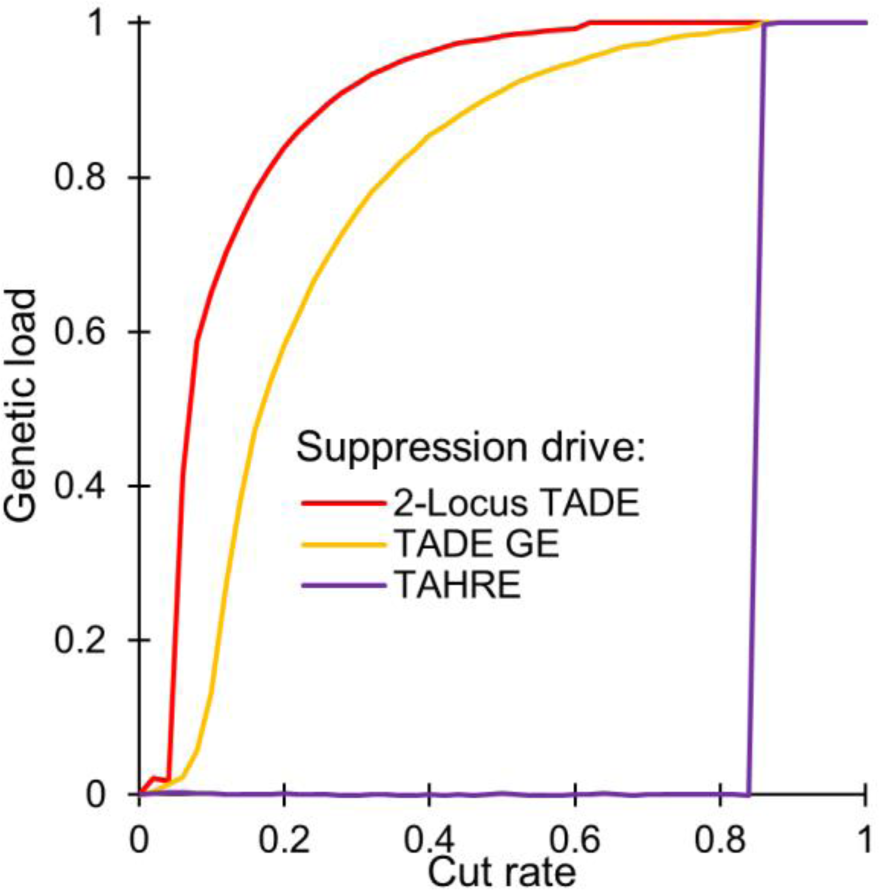
Genetic load of suppression drives. The genetic load imposed on a population as a function of the germline cleavage rate (the embryo cleavage rate does not affect the genetic load) in the suppression drives we considered. We define genetic load as the fractional reduction in the population size of the next generation (caused by the drive at final equilibrium) compared to the expected next generation population size had the population during the present generation been composed entirely of wild-type individuals.

**Figure S2.**
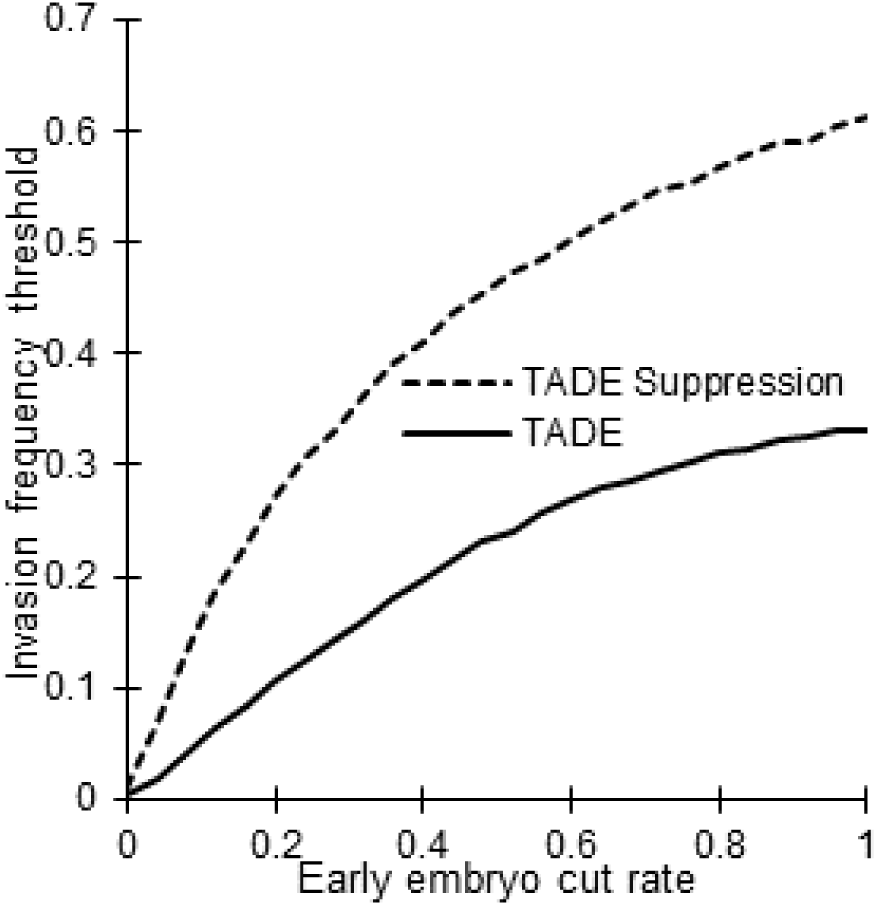
Effect of embryo cut rate on TADE drive thresholds. Invasion threshold frequency in TADE drives as a function of the early embryo cut rate. Released individuals are homozygous for the TADE drive and heterozygous for the TADE suppression drive.

**Figure S3.**
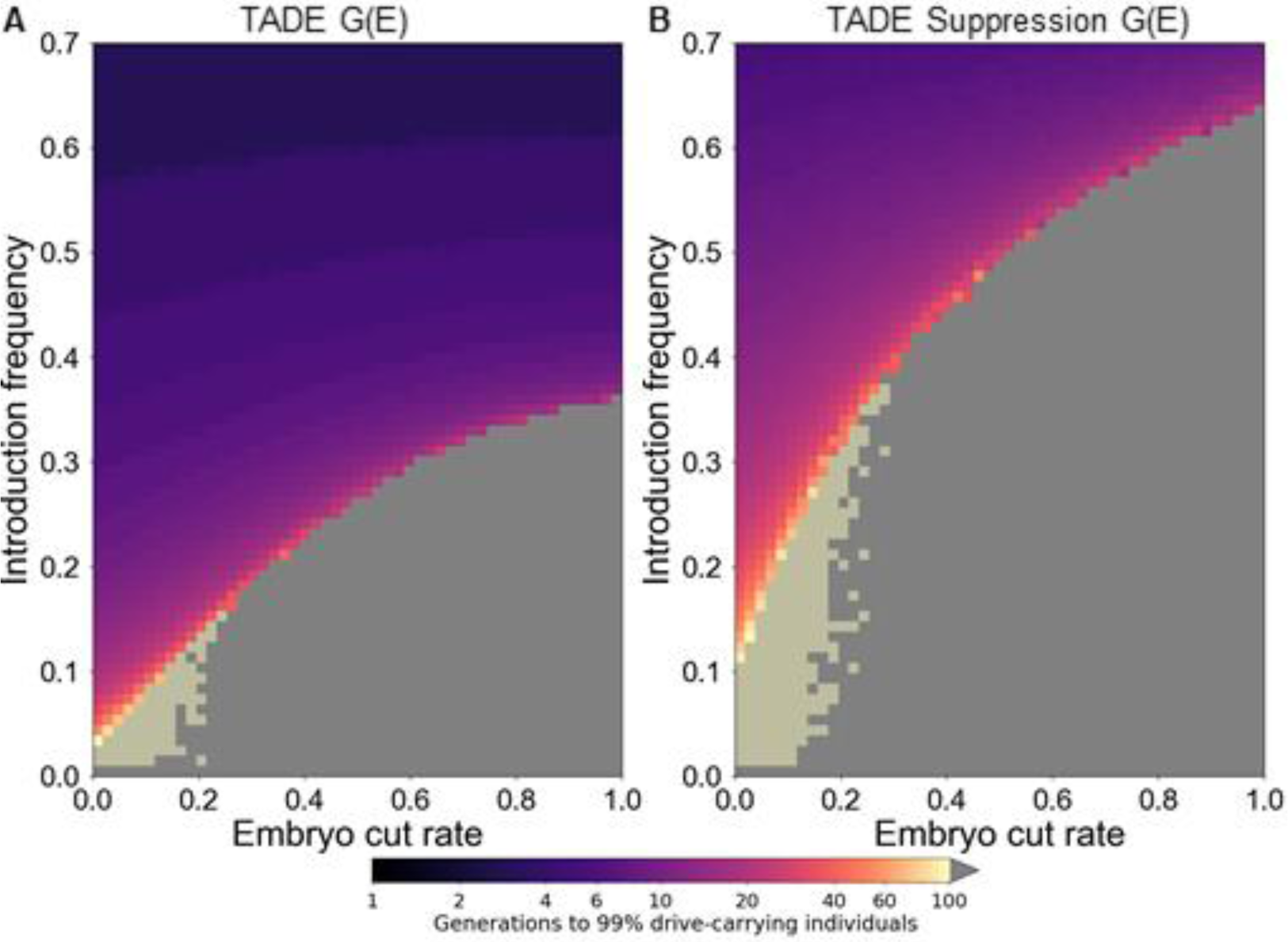
TADE drive performance with variable embryo cut rate. (**A**) The time at which a TADE drive is expected to reach 99% of individuals in the population with varying introduction frequency and embryo cut rate (in the progeny of drive-carrying females). Released individuals are homozygous for the drive allele. (**C**) As in (A), but for a TADE suppression drive (placed in a female fertility gene). Released individuals are heterozygous for the drive allele. Grey indicates that the drive was eliminated within 100 generations. The light sand-colored areas represent regions where the drive does neither reaches 99% of individuals nor is eliminated within 100 generations.

**Figure S4.**
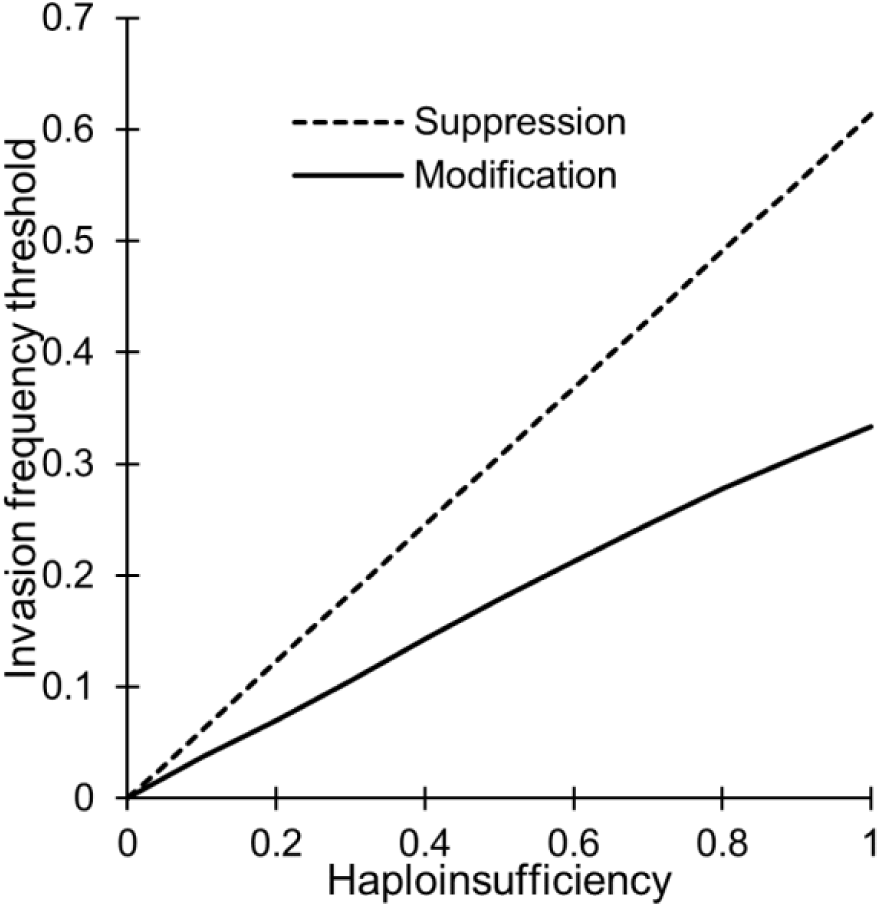
Effect of haploinsufficiency on drive thresholds. Invasion threshold frequency in single-locus TA drives as a function of the degree of target haploinsufficiency. Released individuals are homozygous for the drive allele for the modification drive and heterozygous for the suppression drive.

**Figure S5.**
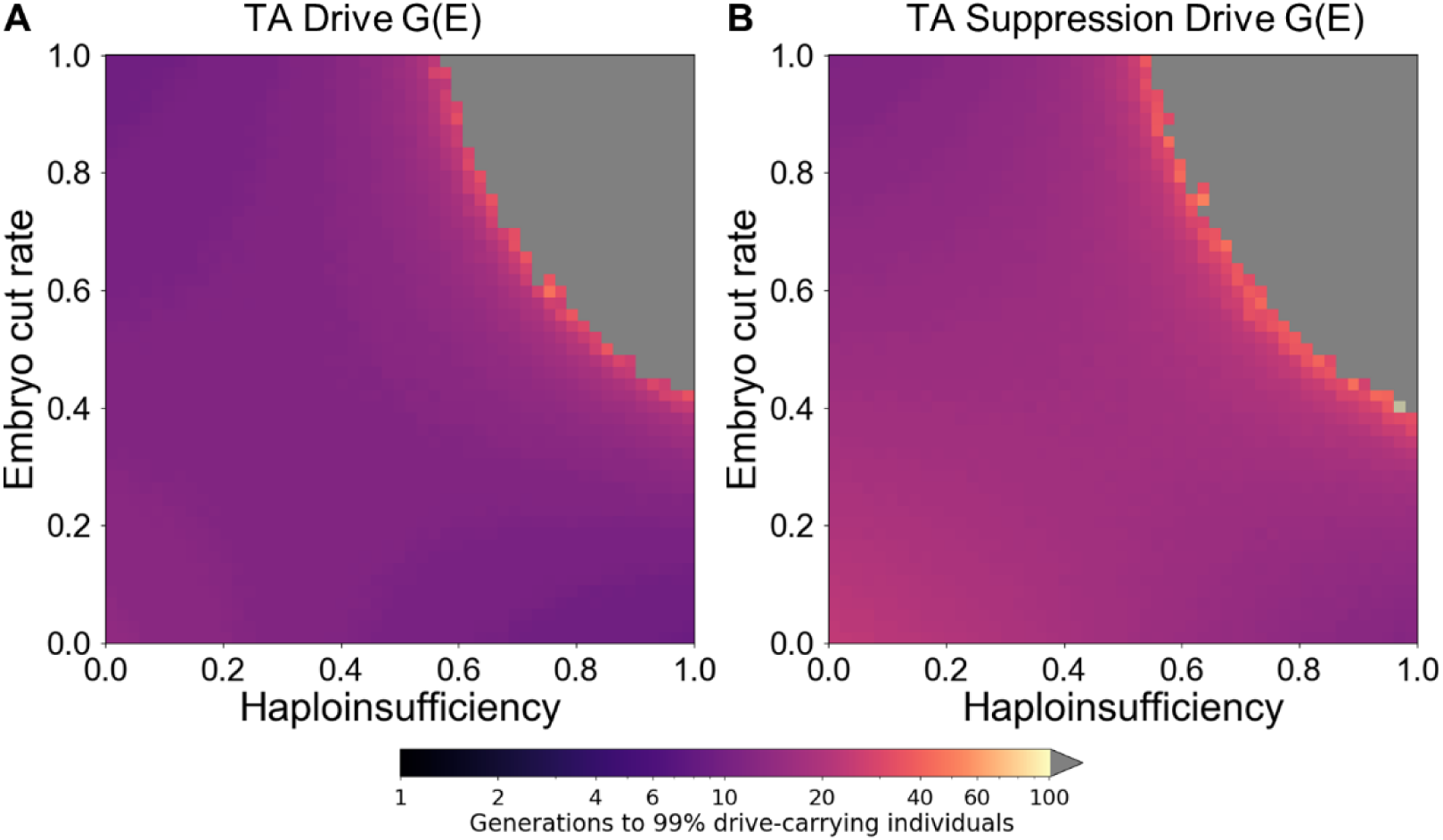
TA drive performance with variable haploinsufficiency and embryo cut rate. (**A**) The time at which a TA drive is expected to reach 99% of individuals in the population with varying haploinsufficiency and embryo cut rate (in the progeny of drive-carrying females). Homozygous individuals are released at 20% initial frequency. (**C**) As in (A), but for a suppression drive (placed in a female fertility gene) and with a release of heterozygous individuals at 40% initial frequency. Grey indicates that the drive was eliminated within 100 generations.

**Figure S6.**
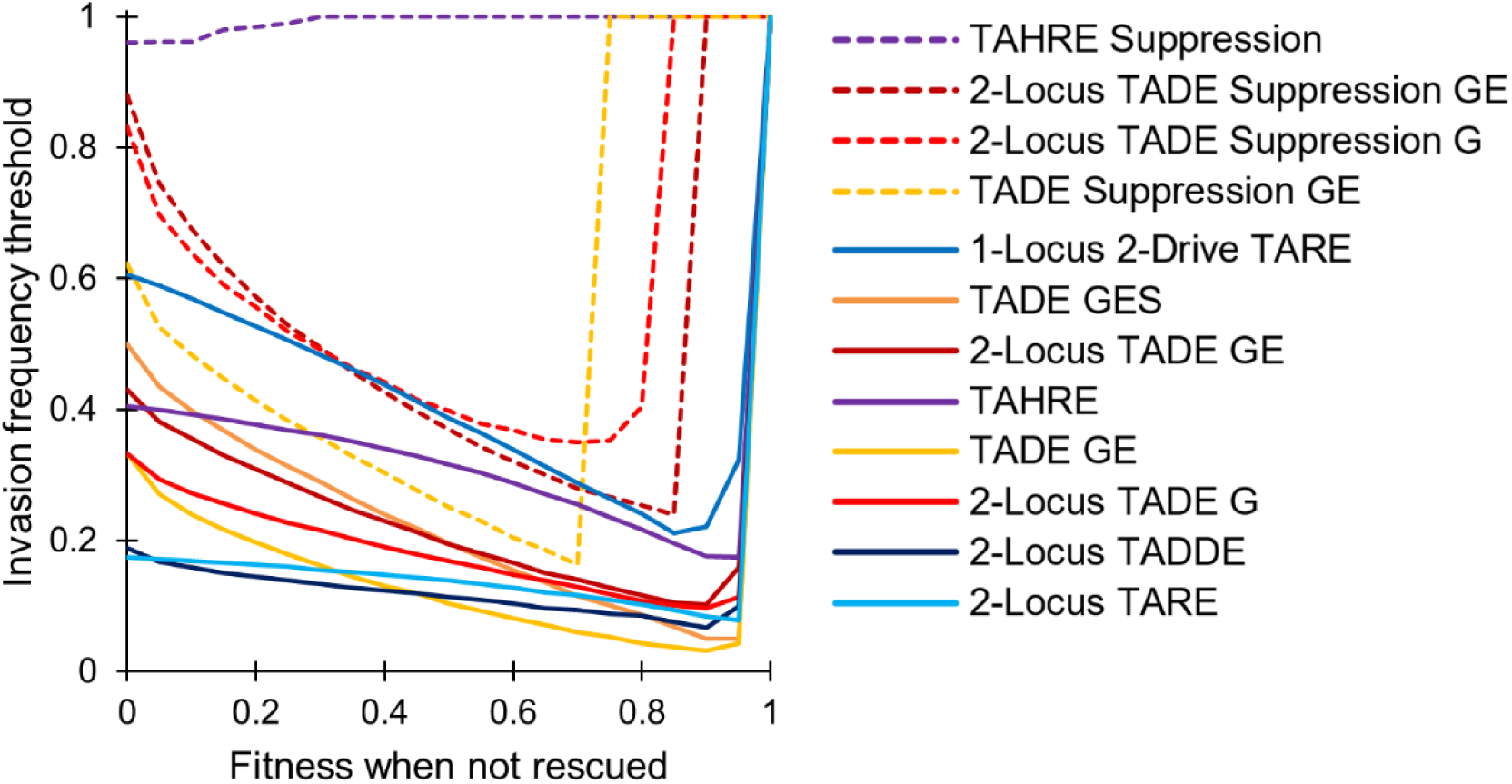
Invasion thresholds of TA underdominance drives with incomplete lethality. Invasion threshold frequencies as a function of the fitness of individuals where rescue is incomplete. In modification systems, released individuals were homozygous for the drive. In suppression systems, individuals were heterozygous for the drive. G = germline only promoter. GE = promoter with germline and early embryo cutting (in the progeny of drive-carrying females). GES = promoter that induces a high rate of somatic cleavage. A threshold of “1” indicates that the system is unable to function as a gene drive.

**Figure S7.**
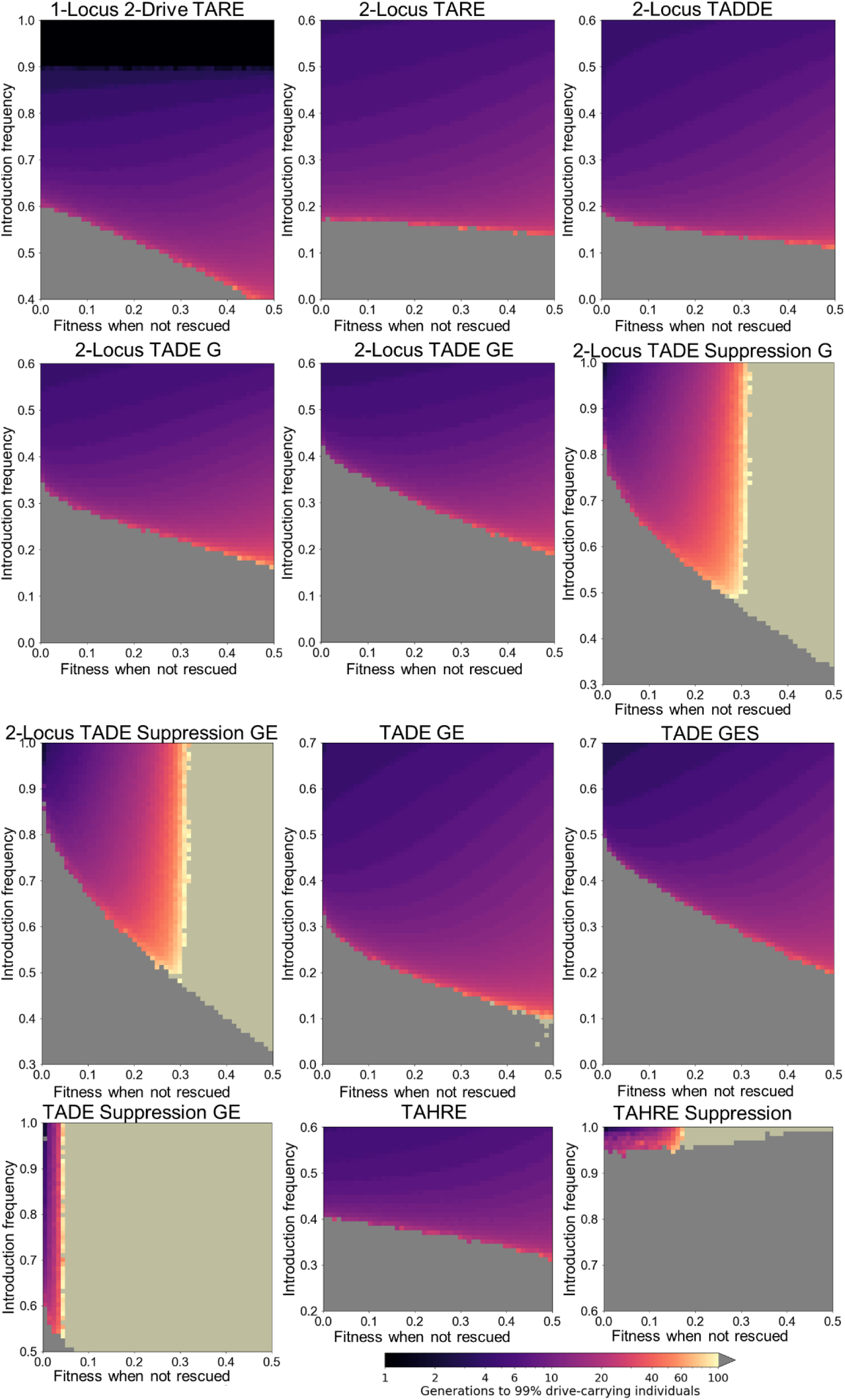
Drive performance with incomplete lethality targets. Heatmaps show the time at which each drive is expected to reach 99% of individuals in the population with varying introduction frequency and embryo cut rate (in the progeny of drive-carrying females). Released individuals are homozygous for the drive allele for modification drives and heterozygous for suppression drives. Grey indicates that the drive was eliminated within 100 generations. The light sand-colored areas represent regions where the drive does neither reaches 99% of individuals nor is eliminated within 100 generations (usually representing a suppression drive that is able to spread and reach a high equilibrium frequency, but that cannot induce a sufficient genetic load to eradicate the population).

**Figure S8.**
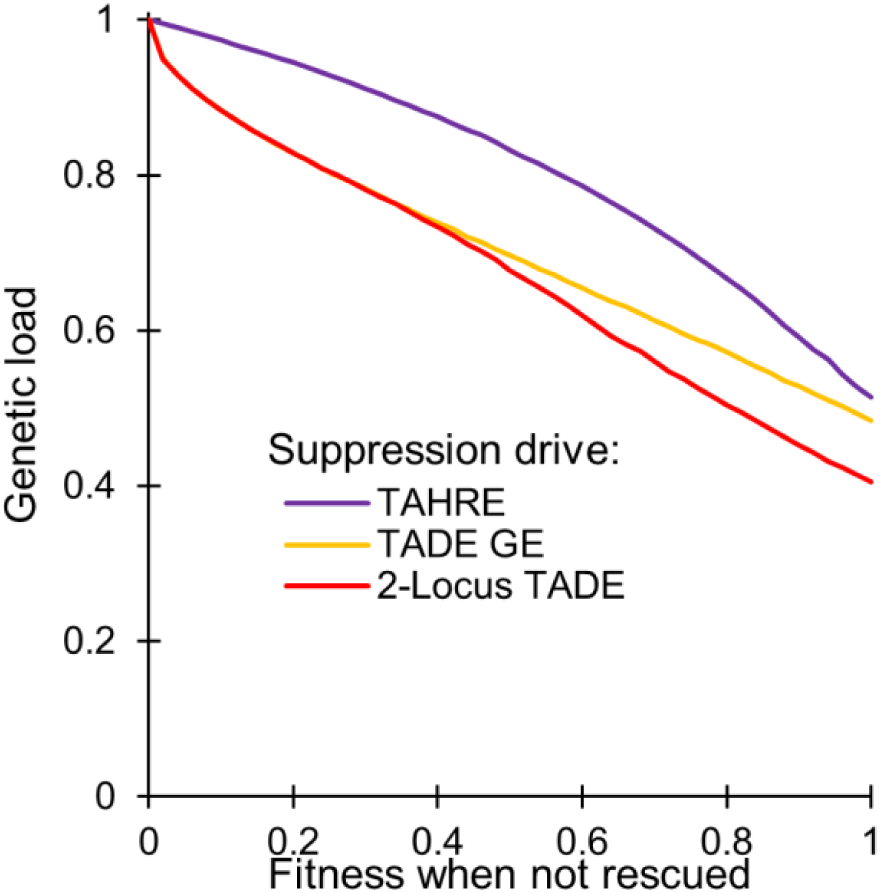
Genetic load of suppression drives with incomplete lethality. The genetic load imposed on a population as a function of the fitness of individuals where rescue is incomplete in the suppression drives we considered. We define genetic load as the fractional reduction in the population size of the next generation (caused by the drive at final equilibrium) compared to the expected next generation population size had the population during the present generation been composed entirely of wild-type individuals.

